# Integration of spatial transcriptomics with immunofluorescence staining reveals spatial heterogeneity and plasticity of astrocytes in experimental glioblastomas

**DOI:** 10.1101/2024.10.11.617740

**Authors:** Mitrajit Ghosh, Paulina Pilanc-Kudlek, Karol Jacek, Szymon Baluszek, Katarzyna Poleszak, Paulina Szadkowska, Bartłomiej Gielniewski, Aleksandra Ellert Miklaszewska, Bozena Kaminska

## Abstract

Astrocytes comprise ∼50% of all brain cells and present distinct morphological, molecular and functional properties in different brain regions. In glioblastoma (GBM), an aggressive primary brain tumour, tumour-associated astrocytes (TAAs) become activated and exhibit different transcriptomic profiles, morphology and functions supporting disease progression. Heterogeneity and specific roles of TAAs within various regions of tumours are poorly known. Advancements of single-cell and spatial transcriptomics allow to profile tumours at unprecedented resolution revealing cell phenotypes, hidden functionalities and spatial architecture in disease-specific context.

We combined spatial transcriptomics and multiple immunofluorescent staining to visualize TAAs heterogeneity and location of various subpopulations in intracranial murine gliomas. Using distinct gene expression profiles, we identified subtypes of TAAs with distinct localization and inferred their specialized functionalities. Gene signatures associated with TAAs reflected their reprograming in the tumour microenvironment (TME), revealed their multiple roles and potential contributing factors shaping the local milieu. Using spatial correlation analysis of the spots, we inferred the interactome of *Slc1a2* (encoding a glutamate transporter) with the other markers of TAAs based on segregated areas of the tumour. The designer RGD peptide blocking tumour-microglia communications, alters the spatial distribution of TAAs in experimental gliomas providing insights into potential mechanisms. Spatial transcriptomics combined with multiple staining unveils multiple functional phenotypes of TAAs and interactions within TME. It shows their distinct morphology and unveils different roles in various regions of the tumour. We demonstrate the glioma-induced heterogeneity of TAAs and their adaption to the pharmacologically-induced modification of the immunosuppressive TME.

## Introduction

Glioblastoma (GBM) is the most deadly, primary brain tumours in adults. The median survival of GBM patients treated with multimodal therapy, including neurosurgery, radiation therapy and chemotherapy, remains only 15 months, and did not change in last 20 years.^1^ The tumour microenvironment (TME) of GBM is heterogeneous in cell composition, highly immunosuppressive and spatially organized. Distinct tumour-associated immune and stromal cells modulate tumour progression, impact therapy outcomes and survival of patients.^2–6^ Astrocytes are most abundant cells in the brain (they comprise ∼50% of all brain cells) and perform critical physiological functions maintaining homeostasis of fluids and neurotransmitters, controlling calcium signalling, synapse maintenance and metabolic supply to support active neurons.^7,8^ The molecular and cellular heterogeneity of astrocytes is poorly defined, mostly by morphological categories as fibrous and protoplasmic astrocytes expressing GFAP (glial fibrillary acidic protein). Combination of transgenic mice, fluorescence-activated cell sorting followed by bulk RNAseq revealed distinct morphological, molecular and functional properties of astrocytes from different brain regions.^9^ Astrocytes are important part of the blood brain-barrier^10–12^, gatekeepers from peripheral insults ^13–16^ and respond to acute insults by undergoing an inflammatory transition to “reactive” state followed by creating a glial scar separating the damaged from the intact brain.^17–19^ A recent high-resolution transcriptomic and spatial cell-type atlas of the whole adult mouse brain was created by combining single-cell RNA sequencing (scRNA-seq) from anatomically defined regions and spatial transcriptomics using multiplexed error-robust fluorescence in situ hybridization (MERFISH). It shows the distinct cell-type organization in different major brain structures and some regions containing cell classes and types highly distinct from the other parts of the brain.^20^ It is likely that astrocytes may exhibit region-specific transcriptional profiles and distinct pattern of activation depending to a disease.

Tumour-associated astrocytes (TAAs) are the abundant cells among GBM stromal compartment and actively interact with GBM cells through diverse cross-talks.^21,22^ “Reactive” TAAs play a crucial role in GBM TME^7,19^, some molecularly defined TAA subpopulations were found in mouse gliomas and analogous populations in primary human brain tumours. ^9^ Two spatially distinct populations of TAAs were identified in mouse gliomas, with those at the tumour periphery being different than those in the perivascular niche. ^23^ TAAs surrounding the tumour resembled morphologically reactive astrocytes, while CD44 and Tenascin-c overexpressing TAAs were restricted to the perivascular niche. Genetic lineage tracing and fate mapping of astrocytes showed that distinct subpopulations expressing GFAP or GLAST (glutamate aspartate transporter) support regional glioma growth in different manner which unravels various functions of TAAs in the spatial context.^24^ The brain with its cellular complexity and spatial organization represents a unique microenvironment, and the mechanisms by which the glioma cells interface with resident populations are not fully defined. While the results of scRNA-seq studies expanded our understanding of gliomas by capturing various malignant cellular states and a vast range of non-malignant cell types in human and experimental gliomas,^25,26^ subclasses of TAAS and their regional localization in GBM are poorly demarcated.

Spatial transcriptomics technologies allow the acquisition of gene expression information from intact tissue sections in the original physiological context at high resolution. Despite some drawbacks such as not achieving single-cell resolution with spot-based spatial transcriptomics or not detecting efficiently some states that are spatially scattered or lowly abundant by unsupervised analysis, ^27^ these techniques accelerated better understanding of the architecture of normal brain and tumour. In this study to dissect the transcriptional diversity of TAAs in a spatial context of mouse gliomas, we combined spatial transcriptomics and immunohistochemistry (IHC) which validated candidate proteins with high cell type specificity. We demonstrate discrete subpopulations of TAAs exhibiting various functional states of tumour-driven activation in spatially resolved locations and provide insights into the relevance of glial scar in malignant gliomas. Moreover, we show that the designer RGD peptide targeting integrin signalling that block reprograming of myeloid cells in the TME and normalizes a vascular niche, affects distribution of TAAs. Altogether, the data provide valuable insights on spatial aspects of transcriptomic heterogeneity of astrocytes in experimental gliomas and the therapeutic modulation of tumour-host interactions that subsequently changes the tumour niche.

## Materials and methods

### Animals

Male C57BL/6J 10-12 weeks mice (Charles River Laboratories, USA) were used for experiments. Mice were housed with free access to food and water, on a 12 h/12 h day and night cycle. All efforts have been made to minimize the number of animals and animals suffering. All research protocols conformed to the Guidelines for the Care and Use of Laboratory Animals (European and national regulations 2010/63/UE September 22, 2010 and Dz. Urz. UE L276/20.10.2010, respectively). Animals were decapitated by a qualified researcher. The First Warsaw Local Ethics Committee for Animal Experimentation approved the study (approval no. 812/2019;1049/2020).

### Glioma cell cultures

GL261 tdT+luc+ glioma cells were generated as previously^28^ described and stably express Firefly Luciferase (luc) and tandem Tomato (tdT) fusion fluorescent protein. GL261 tdT^+^luc^+^ cells were cultured in Dulbecco’s modified Eagle’s medium (DMEM) supplemented with antibiotics (100 U/ml penicillin,100 µg/ml streptomycin) and 10% fetal bovine serum (FBS) (Gibco, MD, USA), Cells were cultured in a humidified atmosphere of CO_2_ /air (5%/95%) at 37°C (Heraeus, Hanau, Germany).

### Stereotactic implantation of glioma cells

Ten weeks old male mice (C57BL/6J) were anesthetized with isoflurane (4–5% induction, 1– 2% maintenance) using Isoflurane vaporizer (Temsega, Tabletop Anaesthesia Station). Before starting the surgical procedure and during the surgery, the depth of anaesthesia was verified by the lack of deep pain response in the limb. Choice of specific anaesthetics was recommended by the veterinarian and approved by The Local Ethics Committee. After identifying the sagittal and coronal sutures on the right side, a hole was drilled at the following coordinates: 1 mm anterior and 2 mm lateral from bregma. GL261 tdT^++^luc^++^ cells (80,000 in 1 μl of DMEM) were stereotactically injected with a Hamilton syringe to the right striatum 3 mm deep from the surface of the brain. The skin was closed and mice were monitored until they completely recovered from anaesthesia. The animals were weighed weekly and observed daily for clinical symptoms. Tumour growth was verified by assaying the luciferase activity of implanted GL261 luc+ /tdT+ glioma cells using Xtreme in vivo bioluminescence imaging system (Bruker, Germany) at 7, 14- and 21-days post-implantation.

### Visium spatial transcriptomics Sample preparation and tissue optimization

Mice were anesthetized and sacrificed by transcardial perfusion with phosphate-buffered saline (PBS). Brains from naïve or tumour bearing mice were removed tumour and snap-frozen in tissue freezing medium (Leica, Ref.14020108926) on dry ice. Brains were coronally sectioned to 10 μm using a cryostat (Thermo Scientific, Microm HM525) at −20°C and mounted onto the etched fiducial frames of the Visium Tissue Optimization Slide according to the Tissue Preparation Guide (CG000240, Rev E, 10X Genomics). A single brain hemisphere section per mouse was mounted on Visium Spatial Gene Expression Slides (catalogue no. 2000233, 10x Genomics). Sections were fixed with pre-chilled methanol at −20°C for 30 min. Haematoxylin and eosin (H&E) staining was performed and subjected to bright-field imaging under a Leica DM4000B microscope according to the staining protocol and imaging guidelines (CG1000160, CG000241, 10X Genomics). Raw images were acquired with a Leica DM4000B microscope and exported as tiff files. Sections were permeabilised with the Permeabilisation Enzyme for different times. The released mRNA was captured by probes on the slides, and reverse transcribed to cDNA marked by fluorescently labelled nucleotides. Tissue was then removed from the slides with a digestive enzyme, leaving the fluorescently labelled cDNA, which was visualized under a Leica DM4000B microscope according to Tissue Optimization Guide (CG000238 Rev E, 10X Genomics). Based on the signal intensity, we determined that the optimal permeabilisation time for a tumour bearing mouse brain with is 26 min. Total RNA was extracted from frozen brains tumour using the RNeasy Kits according to the manufacturer’s instructions (Qiagen). The size, quantity, integrity and purity of all samples were measured using the 2100 Bioanalyzer instrument and RNA chip (Agilent). All sections had an RNA integrity number >8. RNA was eluted in 50 µl RNase-free water and stored at – 80°C until transcriptome profiling.

#### Visium spatial gene expression library construction and sequencing

Visium spatial gene expression slides and reagents kits were used (10X Genomics; #1000187, Dual Index Kit TT Set A; #1000215) and RNeasy Kit (Qiagen; #74134) according to manufacturer instructions (10X Genomics). Sections were fixed in methanol at −20 °C for 30 min and stained for H&E (Sigma-Aldrich; #SLCJ5200, #SLCH5595) for general morphological analyses and spatial alignment of sequencing data. After bright-field imaging, brain sections were enzymatically permeabilised for 24 min, stained and fixed tissue sections were deposited onto the slide. The poly-A mRNA was released from slide covering cells and was captured on each of the spots on the capture area. Each slide spot contained the special barcode composed of an Illumina compatible Truseq sequence, unique barcode, UMI, and poly(dT) sequence. Library preparation was done according to the Visium Spatial Gene Expression User Guide (CG000239, Rev G, 10X Genomics). The concentration of the resulting libraries was determined by a Quantus Fluorometer with a QuantiFluor ONE Double-Stranded DNA System (Promega, Madison, WI, USA), and the quality check was performed using an Agilent Bioanalyzer (Agilent Technologies, Santa Clara, CA, USA). The obtained libraries were pooled together and mixed to achieve sequencing depth recommended by manufacturer instructions based upon the slide coverage. Sequencing was performed on Novaseq 6000 (Illumina, San Diego, CA, USA) with the recommended protocol (read 1: 28 cycles; i7 index read: 10 cycles; i5 index read: 10 cycles; and read 2: 50 cycles), yielding between 150-224 million sequenced reads. The eight dual-index Illumina paired-end libraries were sequenced on a NovaSeq 6000 on an S2 100-cycle flow cell using 150 base pair paired-end dual-indexed set-up to obtain a sequencing depth of ∼50,000 reads as per 10x Genomics recommendations. BCL to FASTQ conversion was performed using SpaceRanger (v1.2.0). Raw FASTQ files alignment to the 10X Genomics mouse reference genome GRCm39–2020 and the reads assignment to spots was performed using SpaceRanger (v1.2.0). The expression matrix was normalized and scaled using the NormalizeData and ScaleData functions from Seurat package^29^ (v4.3.0). Next squared coefficient variance (cv2) modelling was utilized to select most variable genes, principal component analysis (PCA) was run and Marchenko-Pastur algorithm was utilized to identify non-random^30^ components. These data were integrated with canonical component analysis (CCA) in Seurat IntegrateData function. Subsequently, again PCA was run, Marchenko-Pastur algorithm was applied, to uniform manifold approximation and projection (UMAP) was derived, and clustering with Leiden algorithm was performed^31^.

#### Spatial gene expression, profiles and correlation analysis

Differential gene expression between clusters was performed in pairs: tumour core-tumour periphery, tumour periphery - border area, and border area - normal brain. Effect size (log2(fold-change)) calculated between areas using wilcoxauc function from presto package (v1.0.0) was an input for a gene-set enrichment analysis (GSEA) on gene sets from Gene Ontology^31^ and Reactome^32^ databases. The GSEA was performed using the fgsea algorithm from fgsea package (v.1.24.0). Gene pairs correlation was computed using Spearman test within each group of spots (brain, peri-tumoral area, tumour border and core); p-value adjustment was performed with the Benjamini-Hochberg procedure. For comparison of correlation coefficients in each area, Fisher transformation was applied to correlation coefficients and t-distribution was utilized for hypothesis testing (H0: the correlation coefficients are not different, H1: the correlation coefficients are different, α = 0.05).

### Immunofluorescence

The animals were sacrificed 21 days after GL261 tdT^+^luc^+^ cell implantation and perfused with 4% paraformaldehyde in phosphate-buffered saline (PBS). Brains were removed, post-fixed for 48 h in the same fixative solution and placed in 30% sucrose in PBS at 4°C until they sunk to the bottom of the flask. Tissue was frozen in Tissue Freezing Medium (Leica Biosystems, Richmond, IL, USA) and cut in 10 µm coronal sections using a cryostat. The slides were thawed and dried for 5 min at 37°C after being transferred from the −80°C storage. Next, the slides were put in pre-chilled methanol at −20°C for 30 min. The cryosections were washed 3x for 5 min in PBST (0.1% Triton X-100 in PBS), blocked with 3% DS in 0.4% Triton X-100 in PBS for 2 hr at RT, stained overnight with primary antibodies at 4°C followed by 3x washes in PBST and incubation with secondary antibody for 2 hr at RT. All antibodies were diluted in 0.4% Triton X-100/PBS solution containing 3% donkey serum. Sections were washed 3x in PBST for 5 min, once in PBS for 5 min, followed by washing in ultrapure water for 5 min and mounted with Vectashield Vibrance® Antifade Mounting Medium (# ZKO8O3, Vector Labs, US) with or without DAPI (0.001 mg/ml), for counterstaining nuclei. Images were obtained using the Leica DM4000B fluorescent microscope. Following primary antibodies were used for IF staining: anti-GFAP (DAKO; #Z0334), anti-ALDH1L1 (Abcam; #ab177463), anti-S100B (Abcam; #ab52642), anti-Glutamate Transporter (Chemicon; #ab1783), anti-NeuN (Chemicon; #MAP377), anti-TMEM119 (Synaptic Systems; #400004), anti-fibronectin (Merck; #ab2033), anti-laminin (Abcam; #ab11575), anti-transglutaminase2 (Thermo scientific; MA5-12915). Secondary antibodies such as donkey anti-rabbit IgG Alexa Fluor™ 555 (Invitrogen; A31572), donkey anti-rabbit IgG Alexa Fluor™ 488 (Invitrogen; A21206), donkey anti-mouse IgG Alexa Fluor™ 555 (Invitrogen; A31570), donkey anti-rabbit IgG Alexa Fluor™ 488 (Invitrogen; A21202) and goat anti-guinea pig IgG Alexa Fluor™ 488 (Invitrogen; A11073) were used. Primary antibodies were used in 1:50 or 1:100 dilution when secondary antibodies were used in 1:500 dilution. For each staining three different sections from each mouse brain were analysed. In each section 4 ROIs (Region of Interest) were analysed in and around the tumour core.

### Astrocyte cell primary cultures and co-cultures

Primary glial cell cultures were prepared from the cerebral cortices of P0-P2 pups of C57BL/6J mice as previously^6^ described. The cells were suspended in the medium, counted, checked for viability and seeded at a required density in high-glucose DMEM supplemented with Glutamax, 10% fetal bovine serum (ThermoScientific; CA, USA) and antibiotics. Microglia were identified by immunofluorescent staining as Iba1+ cells (>99%), and negative for the astrocyte marker Gfap. The remaining glial cell monolayer was subjected to strong shaking at 180 rpm overnight to remove sparse oligodendrocytes and *>*97% pure astrocyte cultures were^33^ obtained. Primary astrocyte cultures were identified by immunofluorescent staining as >99% GFAP+ and negative for the microglia marker Iba1.

Astrocytes (2.5×10^6^ cells) were seeded in polylysine (PLL)-coated 6-well plates separately in 2 ml of the DMEM Glutamax medium. Astrocyte cultures were co-cultured with glioma cells 48 h after seeding. GL261 tdT^+^luc^+^ glioma cells were seeded on inserts with 0.4-mm pores at density 2.5×10^6^ /insert. After 24 h, the medium was changed to an astrocytic medium, and these inserts were transferred into the plate with astrocyte cells for another 24 hrs. The glass coverslips were used for immunofluorescence or cell lysates were collected for protein analysis by Western Blotting.

### Western blotting and quantification

Whole-cell protein extracts were prepared, resolved by electrophoresis and transferred to a nitrocellulose membrane (GE Healthcare, #10600003). After blocking with 5% non-fat milk in TBST (Tris-buffered solution pH 7.6, 0.01% Tween-20) the membranes were incubated overnight with primary antibody anti-Tgm2 (Invitrogen, #PA5-29356) diluted (1:500) in a TBST with 5% non-fat milk or with anti-GAPDH (Merck, #MAB374) diluted 1:25000 in 5% bovine serum albumin (BSA) in TBST. The primary antibody reaction was followed by 1 h incubation with horseradish peroxidase-conjugated anti-rabbit IgG (Vector, #PI-1000) or anti-mouse (Vector, #PI-2000) which were diluted 1:10000. Immunocomplexes were detected using an enhanced chemiluminescence detection system (ECL) and Chemidoc (Biorad). The molecular weight of proteins was estimated with Cozy prestained protein ladder (High Qu GmbH, #PRL0102c1). Band intensities were measured by densitometry of immunoblots with BioRad Image Lab software.

### Statistical analysis

Each experiment was performed at least three times, on independent passages/cultures, at least in duplicates. Statistical analyses were performed using GraphPad Prism v6.01 (GraphPad Software, Inc., San Diego, CA, USA). The data were plotted as mean ± SD. Differences between the means of the treatments were evaluated using one-way analysis of variance (one-way ANOVA) followed by post hoc Dunnett’s multiple comparison test or one-sided paired sample t-test (p-value) for two groups analysis. P values were calculated using GraphPad software and considered significant when *P < 0.05 (one-way paired t-test).

## Results

### Tumour-associated astrocytes exhibit spatially organized gene expression in experimental gliomas

To investigate the TAAs landscape and explore their distinct spatial location within glioma, we used the 10X Visium spatial transcriptomics platform and immunofluorescence (IF) to validate the results at a single cell level (**Fig. 1A**). In the experiment, 12 fresh frozen mouse brains were analysed, of which 4 samples (2 control and 2 tumour–bearing brains) were used for the Visium and 8 samples (4 control and 4 tumour–bearing brains) for the IF studies. Using spot clustering and spatial mapping, we identified the tumour core from the universal manifold projection (UMAP) and mapped on the tissue section stained with hematoxylin-eosin (H&E) (Fig.1A). To better visualize expression pattern of astrocytic markers in the tumour relative to the surrounding tissue, we identified histologically distinct areas such as the tumour core (TC), tumour periphery (TP), border (BO) and non-tumour brain (BR) on the tissue sections (**Fig. 1B**). We located spots with high expression of well-known astrocyte-specific genes such as *Gfap*, *Aldh1l1* (coding for Aldehyde dehydrogenase 1 family L1, robustly expressed on both body and processes of astrocytes,) and *S100b* (coding for S100-calcium binding protein B, abundantly expressed in astrocytes^16,34^ and mapped them on the tissue section in distinct areas of the brain section (**Fig. 1C**). Cells expressing *Gfap* and *Aldh1l1* localize at the tumor border and form a ring around tumor core consistent with histologically defined “glial scar”, while the expression of S100b is uniformly dispersed in the whole brain section. Using the defined area segregation, we demonstrate distinct expression profiles of *Gfap*, *Aldh1l1* and *S100b* in selected areas shown as violin plots (**Fig. 1D**). This spatial mapping was extended to other marker genes and genes associated with functions of astrocytes. *Rab6a* (coding for Ras-related protein Rab6a, a pan astrocytic marker)^35^ was abundantly expressed, particularly in the cortex, with some expression in the tumour core. In contrast, *Aqp4* (coding for Aquaporin 4; and important water channel marker abundantly expressed in astrocytes^8,16,34^, as well as *Pdpn* (coding for Podoplanin that has been linked with astrogliosis and has expression profile similar as *Gfap*)^36,37^, *Lcn2 (*coding for Lipocalin2) and *Sipr3* (coding for Sphingosine phosphate receptor 3) which are associated with the immunosuppressive activity of astrocytes^38,39^, formed the ring around the tumor (**Fig. 1E**). Several other important marker genes associated with diverse functions of astrocytes such as *Drd1, Spi1, Timp1, Igtp, Cxcl10* clearly demonstrate the spatial heterogeneity in glioma-bearing brains (**Supplementary Figure S1**).

**Figure 1.**
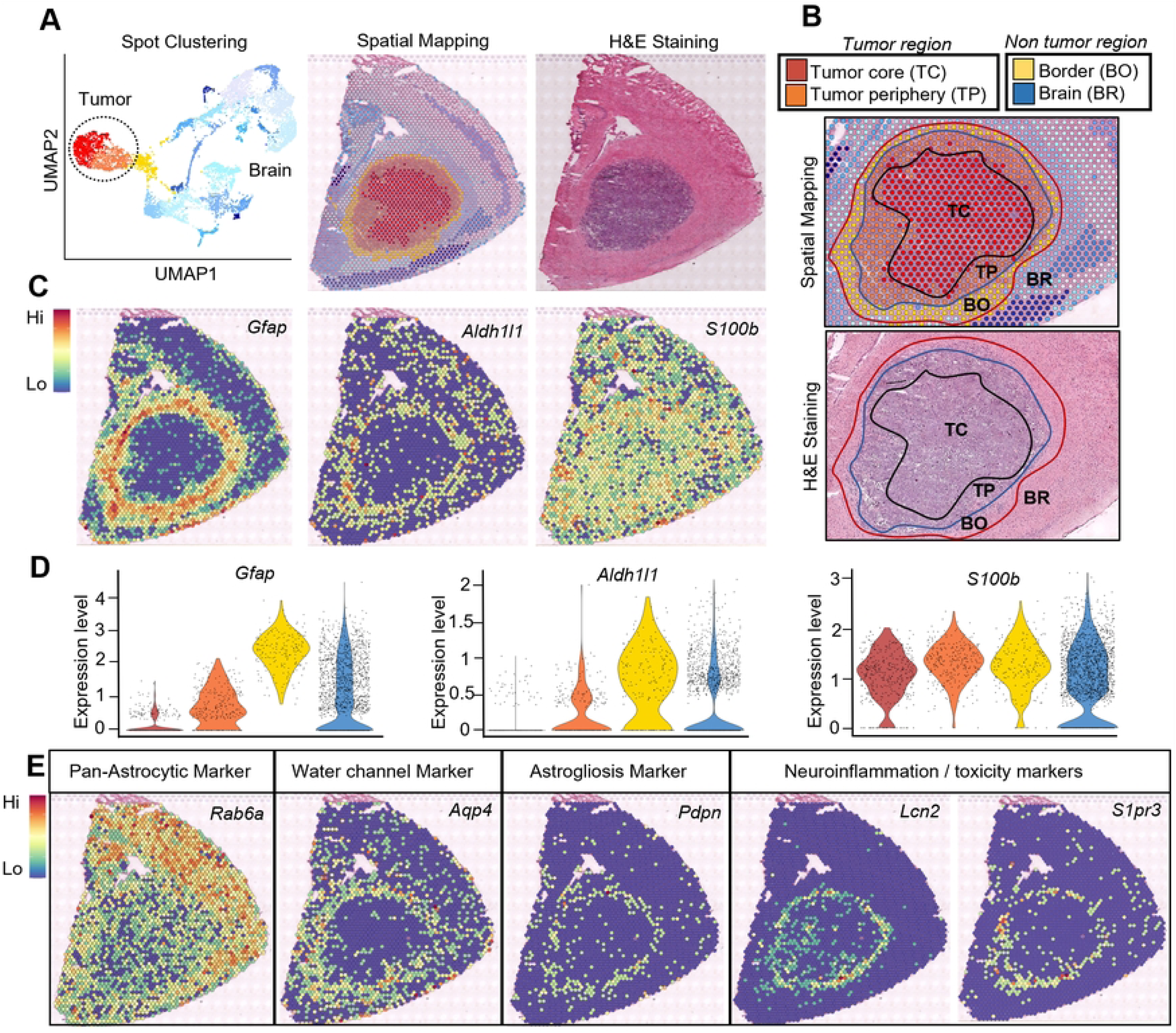
Dissecting the transcriptional heterogeneity of tumour-associated astrocytes in experimental mouse gliomas. (A) UMAP showing brain and tumour cell projection by spot clustering, spatial mapping on the tissue section with the Visium 10X Genomics platform and matching with H&E staining from the same tissue section. (B) Identification and segregation of distinct areas being readouts from UMAP on spatial projection as the tumour core (TC), tumour periphery (TP), border (BO) and brain regions (BR). (C) Spatial mapping of astrocyte marker genes (*Gfap, Aldh1l1* and *S100b*) on tissue sections from Visium gene expression readout. (D) Violin plots showing expression profiles of *Gfap, Aldh1l1* and *S100b* at different tumour and non-tumour regions of the tumour microenvironment. (E) Spatial mapping of *Rab6a* (pan astrocytic marker), *Aqp4* (water channel marker), *Pdpn* (astrogliosis marker), *Lcn2* and *Sipr3* (neuroinflammatory markers).

### Morphology of TAAs varies in different tumour areas

Activation of astrocytes is associated with distinct morphological changes such as hypertrophy, elongation and process extension. Most of an astrocyte surface area is formed by branches, branchlets and leaflets, and a fraction of it can be estimated by glial fibrillary acidic protein (GFAP) immunostaining^8,16^. We performed IF staining for several astrocyte markers on sections of glioma-bearing brains. The pattern of Gfap staining was similar as on the spatial map, with Gfap+ cells surrounding the tumour core and forming a glial scar with the highest expression in the border (BO) area (**Fig. 2A**, upper panel) where the bipolar shaped astrocytes form a thick barrier, with rare cells in the tumour periphery (TP) and almost no Gfap staining at the tumour core (TC) (**Fig. 2A**, lower panel). In the non-tumour parenchyma (BR) Gfap+ cells exhibit the star-like morphology of typical astrocytes. Staining for S100b revealed a similar pattern of morphological differences in all the segmented areas (TC, TP, BO and BR) (**Fig. 2B**). Some S100b+ cells were detected within the TC and the ring in BO was not visible, as S100b+ cells showed more homogeneous distribution. Interestingly, in both cases TAAs close to the BO showed a bipolar morphology aligned along the tumour border, while the cells farther from the BO had mostly star-like morphology (**Fig. 2C**). Double staining for Gfap and S100b verified the presence of TAAs with double positive staining within and around BO (**Fig. 2D**). Aldh1l1 staining shows Aldh1l1+ cells in the BO area surrounding the tumour and rare positive cells within the TC (**Fig. 2E**). Altogether, marker immunostaining confirmed distinct distribution of TAAs characterized by expression of different markers within glioma-bearing brains and corroborated data acquired with the spatial transcriptomics.

**Figure 2.**
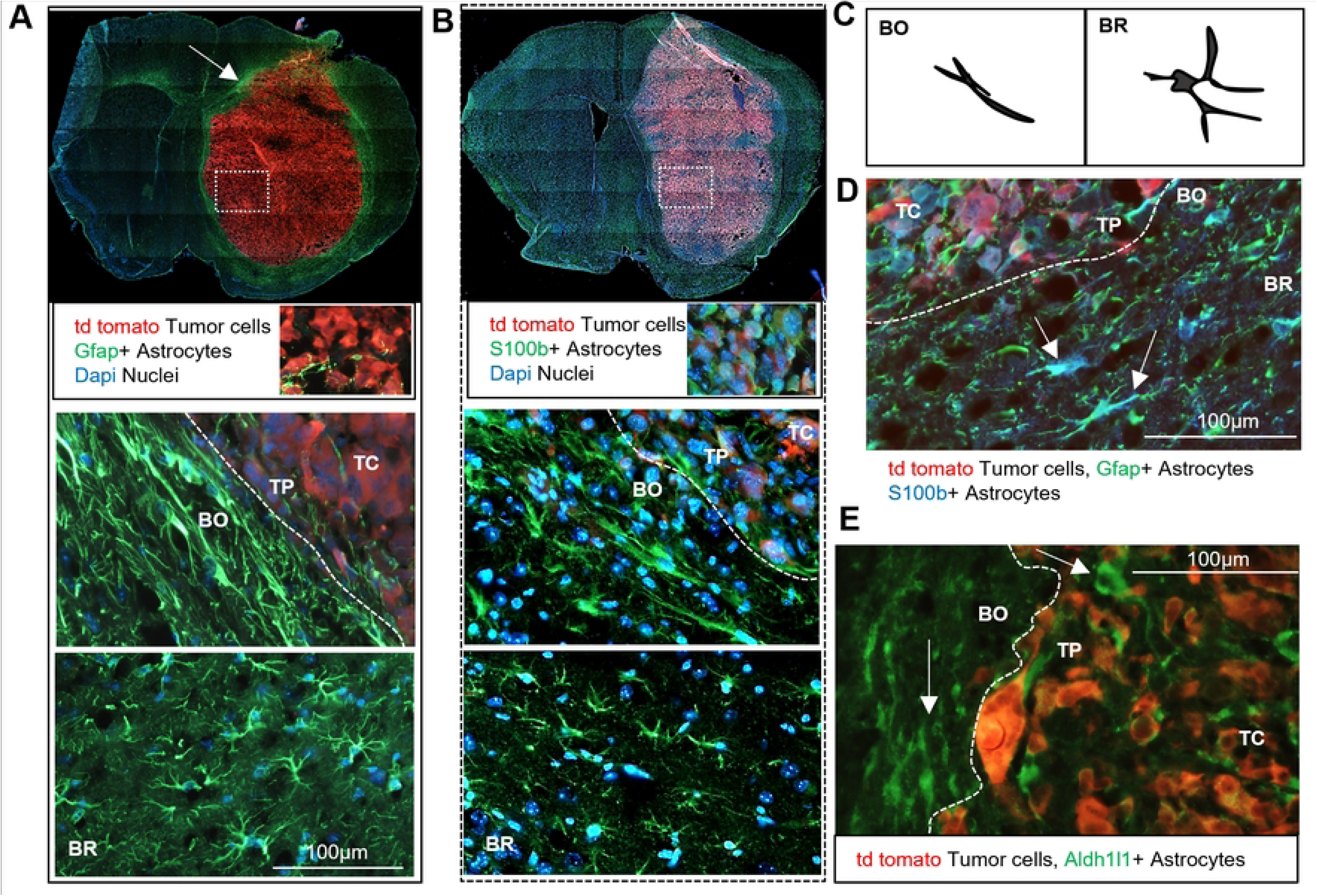
Distinct morphology of astrocytes in various tumour areas. (A) Immunofluorescence (IF) staining of mice brain sections showing td tomato labelled tumour cells (in red), Gfap+ astrocytes (in green) and DAPI stained cell nuclei (in blue) depicts a “glial scar’” composed of reactive astrocytes surrounding the tumour core. Gfap+ astrocytes are barely present in the TC (in the magnified inset), show elongated shapes at the BO and star-like morphology in the surrounding brain parenchyma, BR. (B) IF staining of S100B+ astrocytes (green) shows different patterns, with a uniform, widespread distribution in the brain and the presence of S100B+ astrocytes in the TC and at the BO. S100B+ astrocytes in the BO are elongated, while S100B+ astrocytes farther from the TC display star-like morphology. (C) The scheme illustrates a distinct morphology of astrocytes in BO and BR areas. (D) Double IF staining shows S100B+ and Gfap+ bipolar-extended astrocytes at the BO, whereas in the BR astrocytes display radial-star shapes. (E) IF staining shows Aldh1l1+, bipolar-extended astrocytes (in green) in the TC and BO regions (indicated by arrows).

### Neuron depletion, glutamate dysregulation and astrocyte signatures in the tumour core

In response to pathological conditions astrocytes undergo morphological, molecular, and functional changes, however, there is no prototypical “reactive” astrocyte and divergence binary phenotypes such as good–bad, neurotoxic–neuroprotective, A1–A2, is a useful oversimplification. Reactive astrocytes loss some homeostatic functions and gain some protective or detrimental functions, depending on a context, with only a fraction of common changes between different states. To resolve TAAs functionalities, we sought to integrate the information from multiple staining and spatial data. Staining for NeuN (a neuronal marker) showed a complete lack of NeuN+ cells in the tumour core (**Fig. 3A**) consistent with loss of neurons. Cells expressing neuronal marker genes such as *Rbfox3 (*encoding a marker of post-mitotic neurons RNA-binding FOX3 protein)*, Gria1* (encoding glutamate ionotropic receptor AMPA subunit 1) and *Slc1a2* (encoding solute carrier family 1 member 2 also known as Excitatory amino acid transporter 2 - EAAT2) were absent in the TC and TP, while abundant in the BO and the adjacent BR areas (**Fig. 3B**).

**Figure 3.**
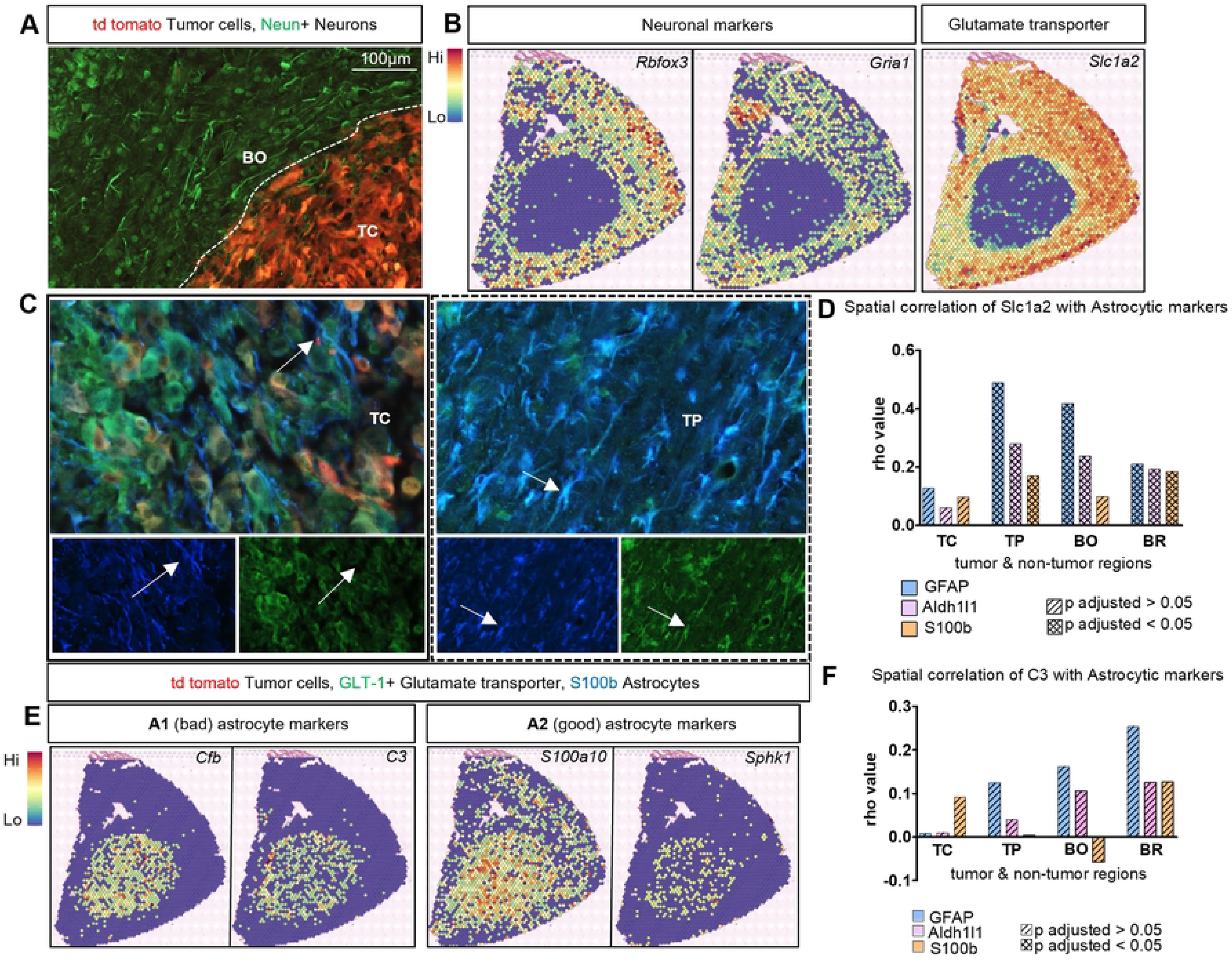
Loss of neurons and glutamate dysregulation coincides with A1 and A2 astrocyte signatures at the tumour core. (A) NeuN+ neurons (in green) are absent in the tumour core demarcated by the presence of td tomato tumour cells (in red), while NeuN+ neurons are distributed uniformly outside of the TC. (B) Spatial mapping of neuronal markers such as *Rbfox3, NeuN*, *Gria1* and glutamate transporter 1 marker *Slcia2/Glt1* in tissue sections confirms a lack of neuronal gene expression and *Slcia2* at the TC. (C) IF staining of GLT-1 (in green) and astrocyte marker S100B (blue) in mouse brain section with td tomato tumour cells (in red). The arrows indicate numerous double positive GLT-1+ and S100B+ astrocytes at the tumour periphery (TP) whereas no double positive cells is visible within the TC; separate channels representing the indicated co-staining. This shows dysregulation of GLT-1 at the TC but not at TP. (D) Spatial correlation extricated from the Visium data indicates stronger correlation between astrocyte genes such as *Gfap, Aldh1l1* and *S100B* with *Slc1a2* at the TP, BO and BR compared to the TC. (E) Spatial mapping of A1 astrocyte specific genes (*Cfb, C3)* and A2 astrocyte specific genes (*S100a10, Sphk1*) on tissue sections from the Visium gene expression readout. (F) Spatial correlation from the Visium gene expression data indicates correlation of cells expressing *C3* with astrocytic markers at different tumour regions.

There are two main mechanisms of neuronal toxicity and death: 1) glutamate dysregulation leading to enhanced excitotoxicity, and 2) neurotoxicity induced by A1 astrocytes emerging under the influence of microglia.^14,19^ The Glutamate transporter 1 (GLT-1) expressed by astrocytes maintains glutamate homeostasis by removing its excess from the extra-synaptic space.^40^ Tumour cells produce an excess of glutamate, however functional astrocytes can maintain its level to support neurons. Cells expressing *Slc1a2* (encoding GLT-1) are less abundant in the TC and TP compared to BO and BR areas on the spatial map (**Fig. 3B**). Double staining for GLT-1 and S100b showed no co-localization of double positive cells in the TC, while such co-localization was observed in the BO area (**Fig. 3C**). This indicates a potential glutamate dysregulation in the TC as TAAs with lower GLT-1 levels do not efficiently remove excess glutamate which may result in neuronal death. Using spatial correlation analysis of the spots, we inferred the interactome of *Slc1a2* with the other markers of TAAs based on segregated areas of the tumour. The graph shows interaction profile of TAAs expressing *Slc1a2* and hints that subtypes of TAAs defined by these marker genes may have different activity profiles based on spatial location in the tumour (**Fig. 3D**).

Under pathological conditions reactive astrocytes acquire distinct phenotypes along the spectrum between the neurotoxic, pro-inflammatory phenotype (A1) and the neuroprotective, anti-inflammatory phenotype (A2). A1 astrocytes are induced by insult-activated microglia that secrete interleukin 1α (Il-1α), tumour necrosis factor (TNF) and complement component 1, subcomponent q (C1q). Their ability to maintain homeostasis and support neuronal survival decreases^14^. Therefore, we investigated the spatial expression of A1 and A2 signature genes such as *Cfb, C3* [A1 markers], and *S100a10, Sphk1* [A2 markers].^14^ Interestingly, all the markers showed clear localization at the tumour site encompassing TC, TP and BO areas and show low expression at non-tumour BR areas; *S100a10* is also expressed in brain parenchyma (**Fig. 3E**). As *C3* represents one of the highly expressed markers for A1 astrocytes, we determined its spatial correlation interactome in the tumour regions where TAAs express *C3* mRNA (**Fig. 3F**), suggesting their neurotoxic capability.

Microglia, brain resident myeloid cells, induce neurotoxic reactive astrocytes and microglia-derived cytokines are the main inducers of the A1 phenotype.^14^ Thus, we determined spatial location of microglia and subpopulations of TAAs in gliomas with ST and IF staining. Tmem119+ non-activated microglia (with radial processes) were found exclusively in the BO and BR regions, whereas the activated microglia (amoeboid shaped) localized mostly at the TP (**Fig. 4A**). Accordingly, *Tmem119* expression was mapped by ST in a proximity to the tumour core and border. Another microglial marker *Sall1*, which is expressed mostly in the non-activated microglia,^41^ shows uniform expression in the brain parenchyma and no expression at the TC (**Fig. 4B**). We demonstrate that genes coding for cytokines and complement: *Tnf, Il1a, and C1qa* are mainly expressed in the TC (**Fig.4C, D**). Therefore, spatial overlap of microglia and cells expressing cytokines, especially at the TC, suggests that these genes are mainly expressed by microglia and influence TAAs, as previous studies with co-cultures demonstrated^14,28^.

**Figure 4.**
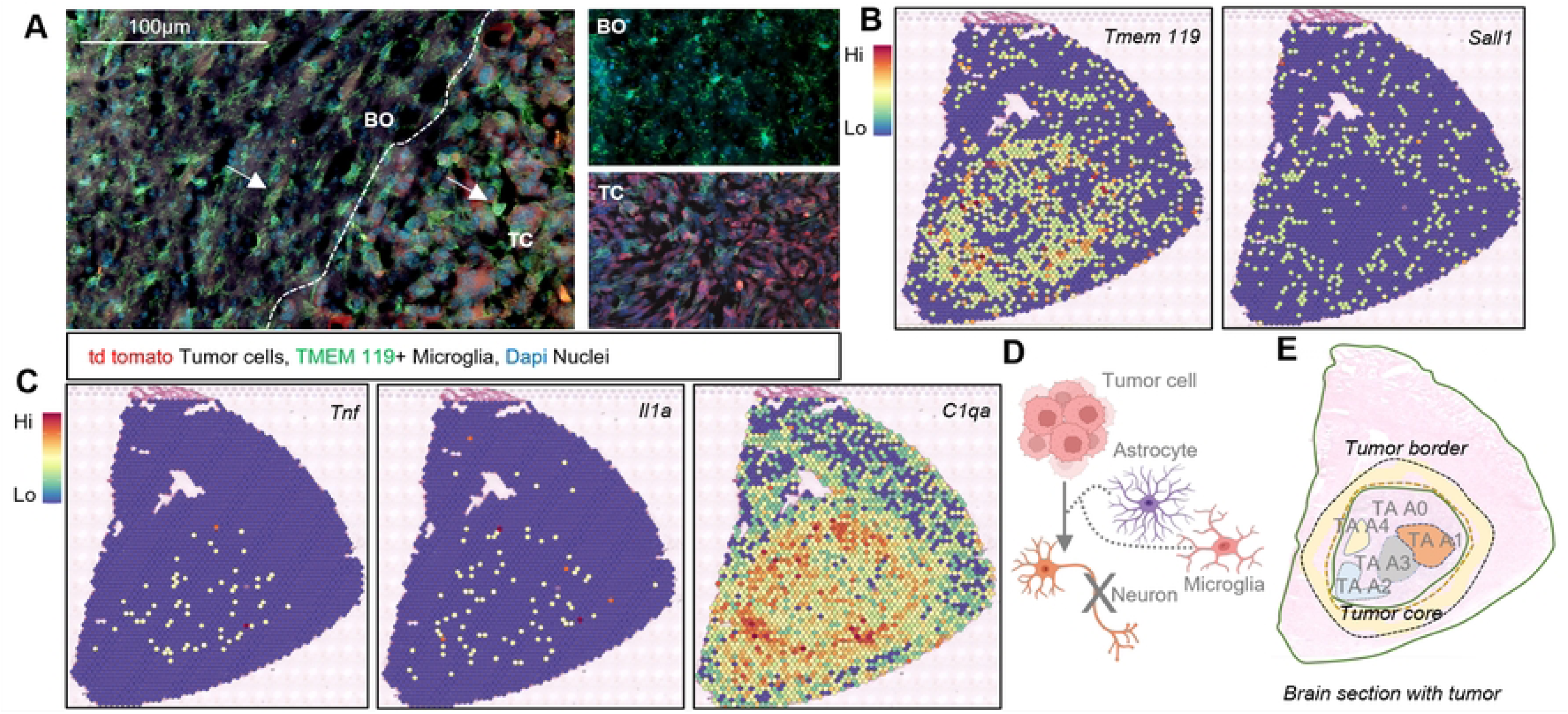
Proximity of microglia influences TAAs in GBM. (A) IF staining reveals TMEM 119+ microglia (in green) with a radial shape (surveying microglia) exclusively in BO and BR regions, whereas microglia with amoeboid shapes (activated microglia) are present at the TC and mostly within TP regions, close to td tomato tumour cells (in red). In the insets; TMEM119+ microglia in BO and TC regions show morphological diversity. (B) Spatial mapping of *TMEM119* expressing microglia in and around the TC, whereas microglia expressing *Sall1* (which is exclusively expressed in surveying microglia) are distributed outside the TC. (C) Spatial mapping of *Tnfα, Il1β*, and *C1qa* depicts localization in the TC. (D) Scheme showing how tumour cells and TAAs influenced by microglia can orchestrate the elimination of neurons at the TC. (E) Schematic representation of at least 5 subtypes of TAAs localized within the tumour -bearing based on marker gene expression depicted by spatial transcriptomics and to some extent validated by immunofluorescence.

Altogether, marker gene distribution, functional correlation and localized occupancy suggest the existence on at least five subtypes of astrocytes in GBM: the A0 subtype (expressing *S100b, Rab6a*) which is homogeneously distributed in the brain, the A1 neurotoxic TAAs expressing *Cfb, C3* and overlapping regionally with cells producing pro-inflammatory cytokines; A2 TAAs expressing *S100a10, Sphk1* markers and located mainly at the TC; A3 *Aldh1l1* expressing TAAs localized in the TC and BO, which has utmost importance for functions beyond Gfap reactivity; A4 the “reactive”, *Gfap* expressing TAAs that form the scar or barrier with high expression at the BO and minimal expression at the TC (**Fig. 4E**).

### TAAs contributes to extracellular matrix remodelling

Composition of extracellular matrix (ECM) in GBM changes during tumour progression due to increased expression of ECM proteins such as laminin and fibronectin, and facilitates diffusive tumour growth.^42^ TAAs have been implicated in ECM remodelling and stiffening the matrix.^43^ We inquired if mechanisms that support diffusive tumour growth show association with the astrocyte heterogeneity. Using Visium ST we mapped cells expressing ECM genes such as *Tenascin C, Laminin b1, Fibronectin 1, Integrin alpha 1,* and found them highly expressed and localized within and around the tumor (**Fig. 5A**). Using IF staining, we visualized increased expression of Fibronectin and Laminin at the TC (**Fig. 5B**) compared to naïve mouse brains without tumour. We queried if there are spatial changes in the expression of ECM related genes that could play a role in stabilization of overexpressed ECM proteins leading to stiffening of the matrix. The tissue transglutaminase or transglutaminase 2 (*Tgm2*) is a Ca^2+^-dependent enzyme that cross-links ECM proteins. Reactive astrocytes produce ECM proteins that become a part of the glial scar and TGM2 overexpressed in those cells may contribute to the ECM protein deposition and aggregation.^44,45^ We found a robust overexpression of *Tgm2* at the TC and TP, while expression of *Tgm1 (*transglutaminase 1) was confined to the BO (**Fig.5C**) where it localized with pro-inflammatory markers such as *Lcn2*, *Sipr3* (described in the Fig.1). Therefore, the ECM transition from a soft to stiff tissue in GBM is likely driven by upregulation of ECM proteins such as Fibronectin, Tenascin C and Tgm2 more likely contributes to their crosslinking (**Fig.5D**).

**Figure 5.**
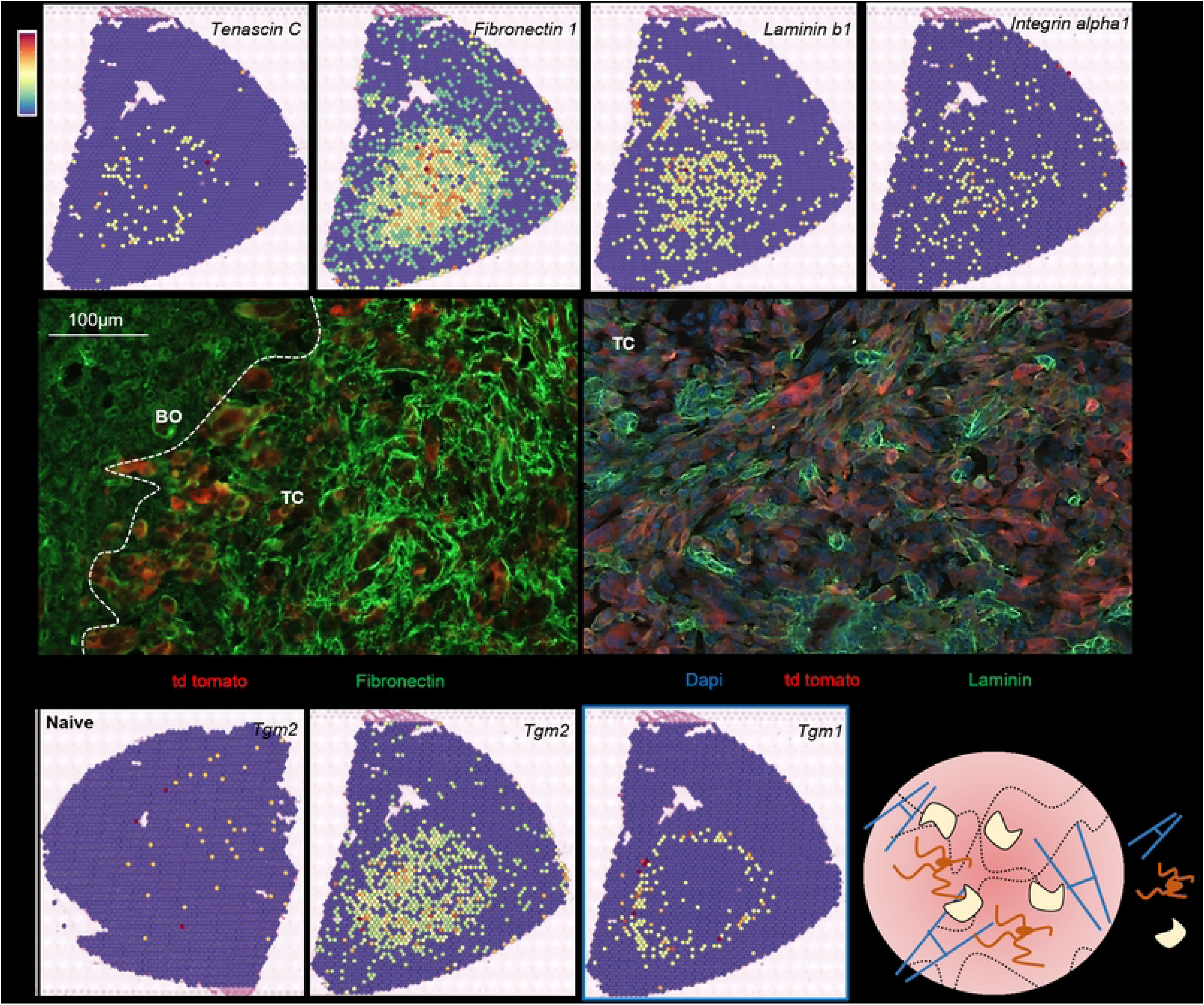
Mapping extracellular matrix remodelling in experimental gliomas. (A) Spatial mapping of *Tenascin C, Fibronectin 1, Laminin b1, Integrin alpha1* encoding important extracellular matrix proteins in sections of brains bearing gliomas denotes localization of these markers in and around TC and TP regions. (B) IF staining shows upregulation of Fibronectin and Laminin in the TC, in agreement with spatial transcriptomics data. (C) Upregulation of *Tgm2* encoding a tissue transglutaminase 2 in sections of brains bearing gliomas whereas there is no expression of *Tgm2* in brains of naïve mice section. Spatial mapping shows that expression of *Tgm1* is restricted to the BO region of the tumour. (D) Schematic representation of extracellular matrix alterations in GBM undergoing from soft to stiff tissue could be regulated by upregulation of ECM proteins such as Tenascin C, Fibronectin, Integrin along with upregulation of a crosslinking protein such Tgm2.

Using primary murine astrocytes and GL261 glioma cells, we examined the impact of co-culture of astrocytes with glioma cells on Tgm2 expression. Tgm2 levels were determined by IF and Western blot analysis (**Fig. 6A**). Tgm2 was detected by IF staining both in primary astrocyte cultures and GL261 glioma cells (**Fig. 6B**), and Tgm2 levels were similar in both cell types. Interestingly, co-culture with astrocytes increased the Tgm2 levels in GL261 cells (**Fig. 6C, D**). It shows that activated TAAs by inducing Tgm2 in GL261 cells may regulate ECM reorganization.

**Figure 6.**
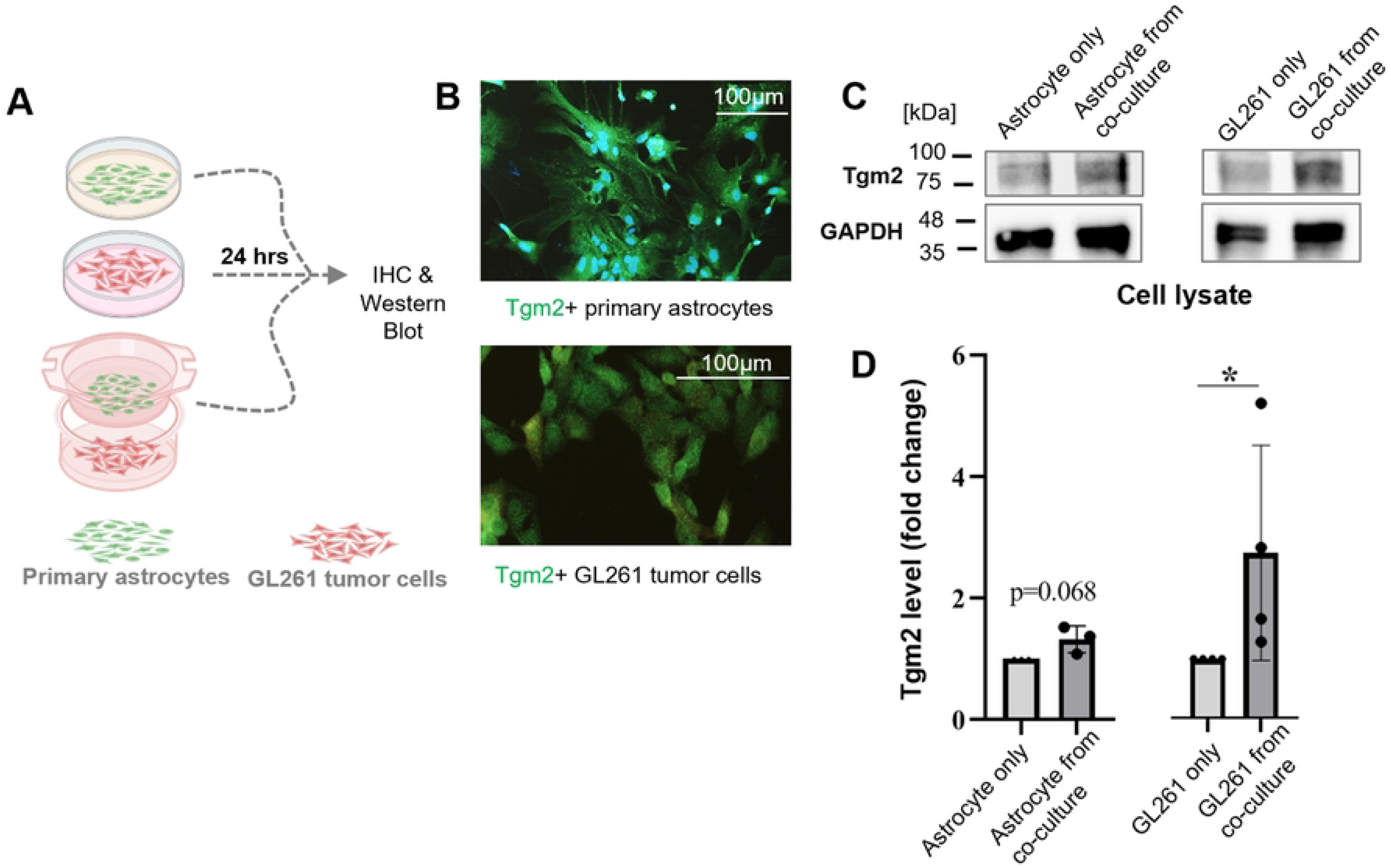
TAAs upregulate Tgm2 in glioma cells. (A) Primary cultures of murine astrocytes and GL261 glioma cells were grown separately or together for 24 h followed by immunohistochemistry and Western blot analysis, n=4. (B) Representative IF images of astrocytes and GL261 cells showing Tgm2 staining. (C) Western blot analysis for Tgm2 from cell lysates of astrocytes, GL261 cells and from those cells growing in co-culture. (D) Quantification of Tgm2 protein levels by densitometry of blots demonstrates increased Tgm2 levels in GL261 cells induced by co-culture with astrocytes. Tgm2 levels were analysed by Western blot and densitometry of immunoblots determined from 4 experiments is represented as mean± SD. Data were normalized to the levels of GAPDH in the same sample; control is set as 1; P values were calculated using GraphPad on logarithmic values and considered significant when *P < 0.05 (one-way paired t-test).

### Blockade of tumour-induced reprograming of myeloid cells reduces astrogliosis in TME and restores GLT1 expression in TAAs

The functional implications of TAAs and formed glial scar in GBM progression are not fully understood. To dissect molecular underpinning of the events, we manipulated the TME using the designer RGD peptide that interferes with glioma-microglia communication and blocks microglia activation via integrin signaling^46,47^. The intratumorally delivered peptide does not affect glioma growth, but blocks the tumour supportive microglia phenotypes and improves immunotherapy with anti-PD1 antibody (unpublished). This allowed to study if a blockade of microglia reprograming, but not tumour growth itself, affects distribution of TAAs. For this purpose, sections from mice at day 21 after tumour implantation and the peptide or control infusion were subjected to Gfap staining. In brains of RGD-treated mice we found a strong reduction of astrogliosis and thickness of the glial scar. Gfap+ astrocytes are significantly more abundant within the TC and reactive Gfap+ astrocytes were reduced at BO and BR areas in the RGD-treated mice compared to controls, which is consistent with the more subdue environment (**Fig. 7A, B**). This shows that tumour-induced activation of astrocytes is likely mediated by activation of myeloid cells in the tumour. The schematic model (**Fig. 7C**) we propose shows there is a spatial heterogeneity of Gfap expression in TAAs in various GBM regions. The change in a density of the astrocyte ring suggests its adjustment to the growing tumour, as it is thicker and denser with GBM progression. Markers of TAAs and morphological structure of the glial scar could be important biomarkers of GBM aggressiveness. Blockade of pro-tumour microglia phenotypes reduced activation of TAAs and restored GLT-1 expression in astrocytes residing in the TC in RGD-treated mice likely limiting neuron death (**Fig. 7D**). Thus, the RGD peptide affecting tumour-microglia communication restores functions of astrocytes which may result in less tumour supporting TME (**Fig. 7E**).

**Figure 7.**
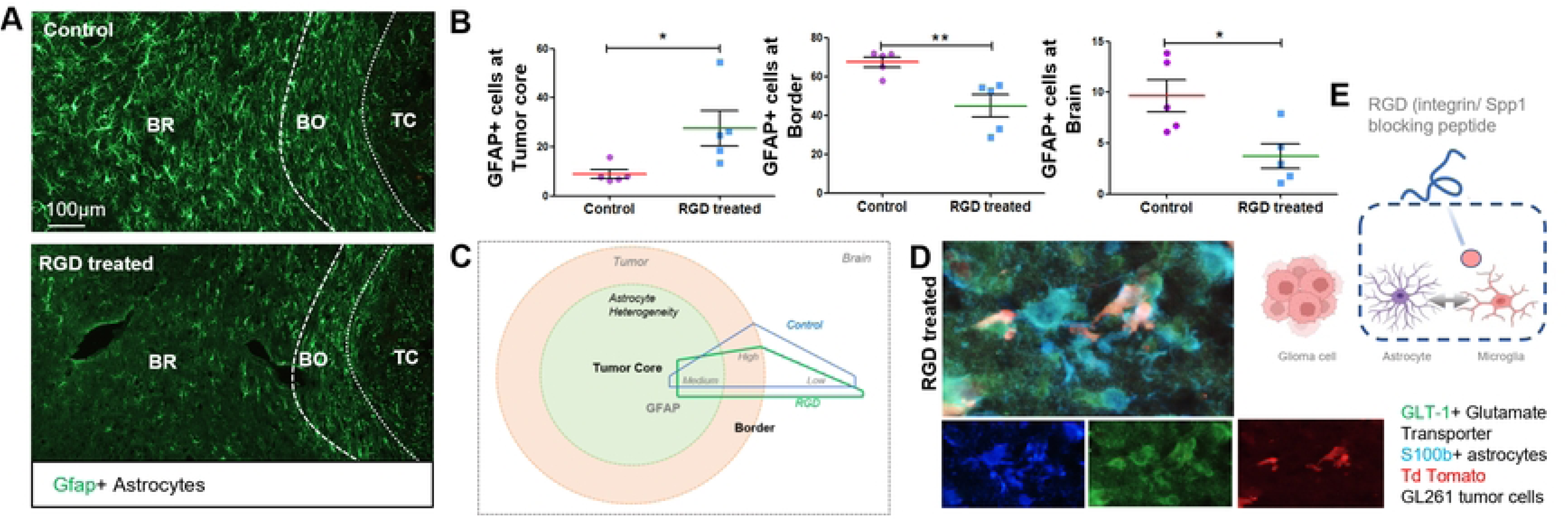
Pharmacologic modification of the tumour microenvironment reduces astrogliosis and harmonizes astrocytic GLT1 expression. (A) In RGD-treated mice less Gfap+, reactive astrocytes are detected in the astrocytic ring around the TC and more Gfap+, reactive astrocytes are present in the TC compared to controls. (B) Quantification of area occupied by Gfap+, reactive astrocytes at TC, BO and BR regions. (C) Schematic representation of astrocytic heterogeneity represented by Gfap+, reactive astrocytes at TC, BO and BR regions and modulation of their distribution in RGD-treated mice. (D) Representative image of GLT-1+ with S100b+ TAAs in brain sections of RGD-treated mice depicts harmonization of GLT-1 expression in S100b+ TAAs. (E) Scheme showing that the blockade of integrin signalling by the RGD peptide impacts microglia functions and disrupts their crosstalk with TAAs, which prevents misusing of those cells in tumour progression.

## Discussion

The identification of diverse astrocyte roles in health and disease led to the concept that there are different astrocyte subtypes that exert different functions. Recent advances in RNA sequencing and proteomics enabled the more detailed assessment of molecular differences between cells and allowed to demonstrate that astrocytes do undergo different morphological and molecular transformations based on age, spatial location and disease context. Dissecting complex cross-talks of TAAs with other cells in various areas of GBM and the role of interactions of a specific TAA subtype were hampered by a lack of distinctive markers beyond GFAP. The data presented here provide the evidence that dynamically changing TAAs play a critical role in shaping TME of GBM and governing tumour progression. We combined 10X Genomics Visium spatial transcriptomics and multiple marker immunostaining to dissect functional phenotypes of TAAs and comprehend the significance of their localization, as well as to understand a role of specific astrocyte subtypes in the local milieu in GBM. Using specific gene marker profiles (in addition to a well-known marker *Gfap*) we found five distinct transcriptional subtypes of astrocytes based on markers expressed in diverse tumour regions, their distance from the tumour core and unique morphology. These transcriptomic profiles have implications for distinct functions of TAAs in tumour progression. We detected astrocytes expressing classical marker genes (e.g., *Aldh1l1*, *Gfap*, *Aqp4* and *Slc1a2*) in control brains and their abundance in tumour-bearing hemispheres where they formed a glial scar around the tumour core. There was a striking difference between bipolar shape of Gfap+ TAAs along the tumour core and star-like morphology of astrocytes that were located farther for the tumour core. We demonstrate the maintenance of regional heterogeneity by TAAs in GBM, which adds an additional layer to the known astrocyte heterogeneity described in brain development, aging and inflammation.^16,34,48,49^ Specifically, we found “reactive” TAAs (positive for Gfap) but also *Aldh1l1* expressing TAAs that potentially contribute to glial scar formation, glutamate dysfunction, neurotoxic influence and ECM reorganization via regulation of Tgm2. Aldh1l1 is one of the most reliable and important markers for astrocytes, though its expression did not change in reactive insults.^16,34^ A recent study of spatial organization of human GBMs defined the reactive astrocyte cluster characterized by classical astrocytic markers (*AGT* and *GJA1*) and genes coding for metallothionein.^50^ Injury-induced glial scar serves as a physical and chemical barrier to axonal regeneration as its major constituents (e.g., Tenascin, N-sulphated heparan sulphate proteoglycans, chondroitin sulfate proteoglycans and keratan sulphate proteoglycans are inhibitory for neurite outgrowth.^15,17^ The glial scar *in vivo* may differ in distribution and type resulting in different biological effects.

We found Aldh1l1 expressing, A1 neurotoxic astrocytes to be determinants for regional stratification at the TC of GBM. The spatial data strongly suggests a proximity of microglia in and around the tumour and its capacity to produce pro-inflammatory cytokines can regulate the behaviour of astrocytes. We observed changes in expression of genes coding for several ECM proteins (Fibronectin, Laminin B1) in the TC along with the increased expression of *Tgm2*. Upregulated levels of those proteins in the TC revealed by IF demonstrate a robust remodelling of the tumour core. The localized expression of *Tgm2* in the TC suggests its role in the reinforcing the ECM re-organization. Tgm2 crosslinks various ECM proteins, including fibronectin, fibrinogen/fibrin, von Willebrand factor, vitronectin, dermatan sulfate proteoglycans, collagen V, osteonectin, laminin, and osteopontin. Astrocytic Tgm2 has been shown to facilitate cell migration and proliferation, reducing their ability to protect neurons after brain injury. Moreover, Tgm2 supports glioma stemness and radioresistance.^45^ We found that glioma cells increased levels of Tgm2 in the presence of TAAs. As Tgm2 regulates ECM stiffening by altering the composition of ECM proteins, these properties make the enzyme an important target for GBM therapy.

We explored if glioma-induced changes in microglia functions affect astrogliosis in gliomas independently of glioma growth. Therefore, we employed a designer peptide RGD that specifically blocks microglia-glioma communication and reverses microglia reprogramming in TME when delivered intra-tumorally.^46,47^ We show that TAAs are modulated as a consequence of the treatment and changed their morphology at the glial scar. The glial scar forms a physical barrier for cell-cell communication, cell infiltration or even for drug penetration.^19,49,51,52^ We found that in RGD treated tumours the glial scar is less dense with more astrocytes within the TC and those TAAs express GLT-1 that suggests a normalizing effect on TME.

Spot-based spatial transcriptomics employed in this study has its limitations such as low resolution, making resolving individual cell states and cell types challenging. However, a recent study showed that cells tend to be surrounded by other cells in the same state forming local environments, highly enriched with cells in an individual state. While the 10X Visium technology does not provide single-cell resolution, as a diameter of a spot is around 55 μm, they found a spot contains a mixture of 1–35 cells, with a median of 8 cells in GBM based on image analysis.^27^ Despite this limitation, the transcriptomic patterns detected in this study provide meaningful assumptions and the findings were confirmed by multiple staining for astrocytic markers. Altogether, by combining spatial transcriptomics and immunostainings we resolved astrocytic heterogeneity, demonstrated astrocyte regulation within TME and its impact on glial scar and tumour expansion. The experiments with the RGD peptide modifying TME allowed to dissect TAAs roles and their impact on other cells more precisely.

## Data availability

All data supporting the findings of this study are available within the manuscript or the supplementary information or from the corresponding author upon a reasonable request. The raw and processed data are available in GEO under accession number GSE269545.

## Acknowledgements

We would like to thank Ms. Beata Kaza for technical help and sharing her expertise.

## Funding

M.G. is supported by PACIFIC Call 1 (PAN.BFB. S.BDN.619.022.2021, the EU Horizon 2020 research and innovation programme under the Marie Skłodowska-Curie grant 847639). Studies were supported by grant 2020/39/B/NZ4/02683 from the Polish National Science Center (BK).

## Declaration of competing interests

The authors declare no competing interests.

## Abbreviations

GBM: glioblastoma
GFAP: glial fibrillary acidic proteins
DMEM: Dulbecco’s Modified Eagle Medium
TAAs: tumour-associated astrocytes
TME: the tumour microenvironment.

## Authors’ contributions

MG conceived the idea, performed the experiments, analyzed and interpreted the data, and wrote the manuscript; PPK & KP performed some experiments; KJ & SB performed all the bioinformatic analysis; PS & BG performed the sequencing; AEM analyzed and interpreted the data; BK conceived the idea, obtained funding, analyzed and interpreted the data and wrote the manuscript. All authors read and approved the final manuscript.

## Notes

### Competing Interest Statement

The authors have declared no competing interest.

